# SSDraw: software for generating comparative protein secondary structure diagrams

**DOI:** 10.1101/2023.08.25.554905

**Authors:** Ethan A. Chen, Lauren L. Porter

## Abstract

The program SSDraw generates publication-quality protein secondary structure diagrams from three-dimensional protein structures. To depict relationships between secondary structure and other protein features, diagrams can be colored by conservation score, B-factor, or custom scoring. Diagrams of homologous proteins can be registered according to an input multiple sequence alignment. Linear visualization allows the user to stack registered diagrams, facilitating comparison of secondary structure and other properties among homologous proteins. SSDraw can be used to compare secondary structures of homologous proteins with both conserved and divergent folds. It can also generate one secondary structure diagram from an input protein structure of interest. The source code can be downloaded (https://github.com/ethanchen1301/SSDraw) and run locally for rapid structure generation, while a Google Colab notebook allows easy use.

## Introduction

Recent advancements in cryo-electron microscopy^1^, metagenomics^2; 3^, and deep learning-based protein structure prediction methods^4-7^ have led to an explosion in the number of available protein structures. For instance, the number of entries in the Protein Data Bank^8;9^ (PDB), a repository of experimentally determined protein structures, has nearly doubled in the past 10 years. Earlier this year, the authors of ESMfold–a large language model that rapidly predicts three-dimensional protein structures from single sequences–released a web-based collection of >772 million predicted structures predicted from metagenomic sequences^6^. Similarly, >200 million structures predicted by AlphaFold2^4^–a highly accurate deep-learning based model for protein structure prediction–are through a web repository for easy user access^10^, and many have been deposited into the UniProt sequence database^11^.

The enormous increase in available models of protein structure presents opportunities to identify large-scale relationships between structure and other properties, such as sequence conservation or prediction confidence. Such relationships are often most effectively depicted when multiple protein structures are compared, motivating the development of structural alignment algorithms that match common elements of protein structure rather than amino acid sequence^12^. Nevertheless, important relationships between protein structures can be obscured by three-dimensional visualizations that cannot effectively convey all structural features through one image. This shortcoming especially impacts homologous proteins with non-conserved structural features arising from insertions, deletions, or mutations that cause substantial changes in secondary structure. Indeed, the need for easily interpretable structure diagrams is underscored by several recent studies highlighting how protein structure can transform dramatically in response to seemingly minor sequence changes^13-17^. To observe these transformations accurately, secondary structures of the proteins of interest must be registered, meaning that amino acids with annotated secondary structures must be aligned with their corresponding amino acids in a multiple sequence alignment (MSA). Once registered, secondary structure structures of homologous proteins aligned within the MSA can be compared, and their respective secondary structure diagrams become comparative. That is, the secondary structure of Protein A at position X can be compared directly to the secondary structure of its homolog, Protein B, at position X if their secondary structures are both registered to the same MSA (**Figure 1**). Comparative secondary structure diagrams also simplify the visualization of fold-switching proteins, single sequences evolutionarily selected to remodel their secondary and tertiary structures in response to cellular stimuli^18-20^. In short, as increasing evidence indicates that highly similar or identical protein sequences can assume folds with drastically different secondary structures^21^, the need to graphically depict structural differences among homologous proteins and relate them to other protein properties increases.

**Figure 1.**
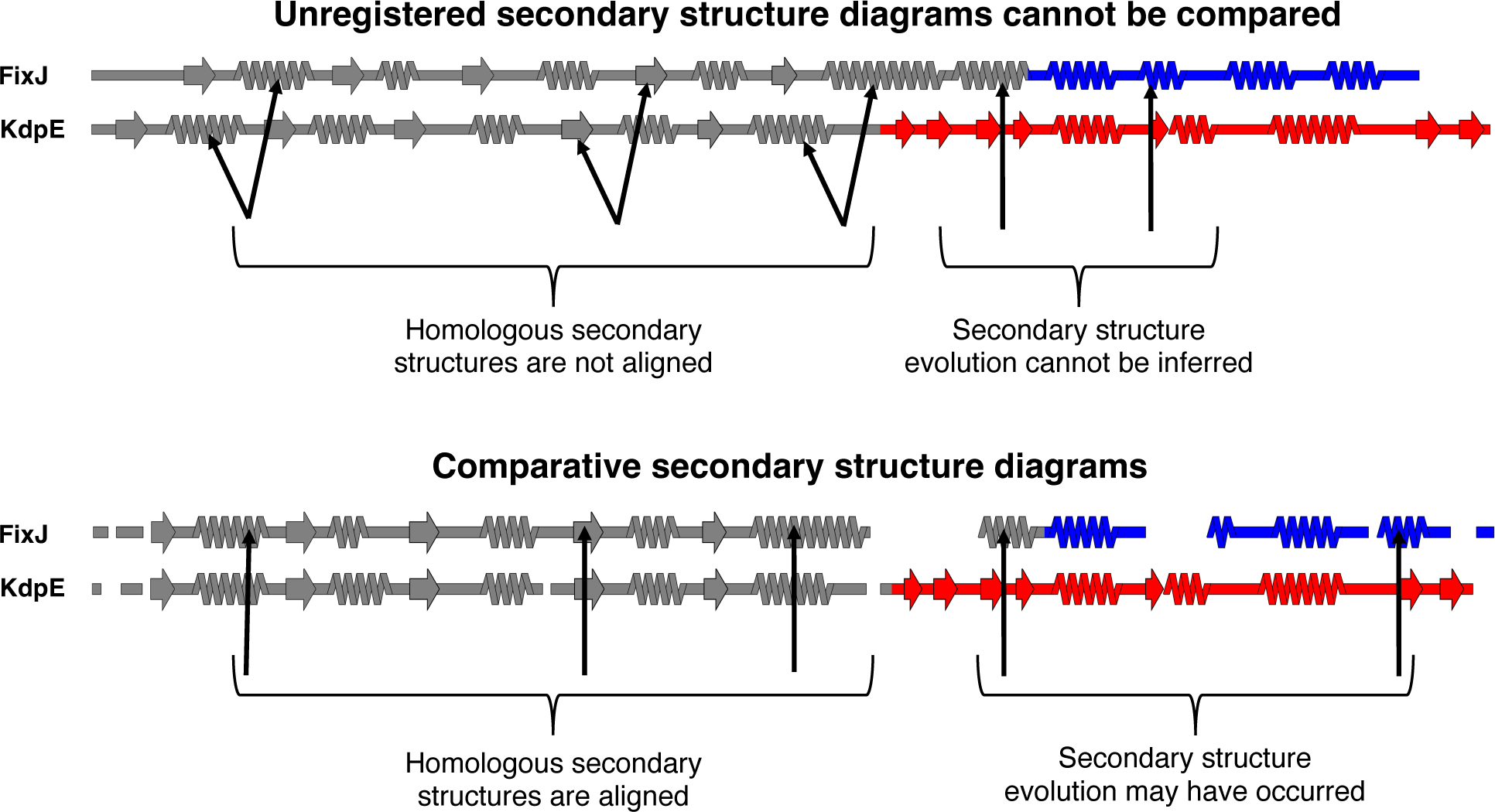
Comparative secondary structure diagrams result from registering secondary structure annotations with their corresponding aligned amino acid sequences. Secondary structures of unregistered diagrams (above) are not aligned, disallowing reliable inferences about secondary structure evolution. Diagrams align when their secondary structures have been registered with their corresponding aligned amino acid sequences (comparative secondary structure diagrams, below), suggesting possible secondary structure evolution where α-helices align with β-sheets. Secondary structure diagrams were made from structures of bacterial response regulators FixJ (PDB ID: 5XSO, chain A) and KdpE (PDB ID: 4KFC, chain A). The C-terminal domains of FixJ and KdpE are colored blue and red, respectively, indicating different folds (helix-turn-helix, blue; winged helix, red), while their structurally conserved N-terminal domains are gray. Arrows pointing to the gray domains indicate homologous secondary structures; arrows pointing to the colored domains indicate divergent secondary structures. Previous phylogenetic analysis and ancestral reconstruction^17^ indicate that the C-terminal β-sheet of the winged helix evolved from the C-terminal α-helix of the helix-turn-helix.

To effectively depict relationships between the structures of homologous proteins and other properties of interest, we present SSDraw, a Python-based program that rapidly generates secondary structure diagrams from three-dimensional protein coordinates. These linear diagrams are registered to a user-inputted MSA and colored by any property of interest. Running SSDraw once generates a diagram of one protein from an MSA. Multiple diagrams from one MSA can be generated and stacked for easy comparison. These functionalities distinguish SSDraw images from other secondary structure visualizations^22-28^. For instance, ESPript^23^ relates secondary structures derived from one representative protein structure to multiple homologous sequences, usually divided on multiple lines of text. This format works well when the user seeks to visualize sequence conservation patterns in a protein family with conserved secondary structures. SSDraw may be preferable if the user seeks to compare structures of homologous proteins with divergent secondary structures by stacking each diagram and comparing structural differences. As another example, secondary structure diagrams from Aquaria^25^ also generate stackable linear secondary structure diagrams but color by sequence conservation only. SSDraw may be preferable if the user seeks to color the stacked diagrams by a property other than sequence conservation. In short, SSDraw was written to fiexibly relate secondary structure differences between homologous proteins with other protein properties of interest. While this software was originally designed for fold-switching proteins^19^ and homologous sequences that with different secondary structures^17^, it also serves as a tool to quickly generate secondary structure diagrams for individual proteins with custom coloring by sequence position in seconds (local install) to minutes (Google Colab notebook).

## Results

### Software overview

SSDraw requires two inputs to run: (1) a file containing three-dimensional protein coordinates in PDB format and (2) a multiple sequence alignment in FASTA format (**Figure 2**). SSDraw requires only alpha carbon coordinates to generate an image. The user may specify the chain ID if they input a multi-chain PDB. The multiple sequence alignment can be generated with programs such as MUSCLE^29^, Clustal Omega^30^, or HMMER^31^, so long as it is inputted in FASTA format. The user may also input a single ungapped FASTA sequence if they are interested in generating a diagram from a single sequence.

**Figure 2.**
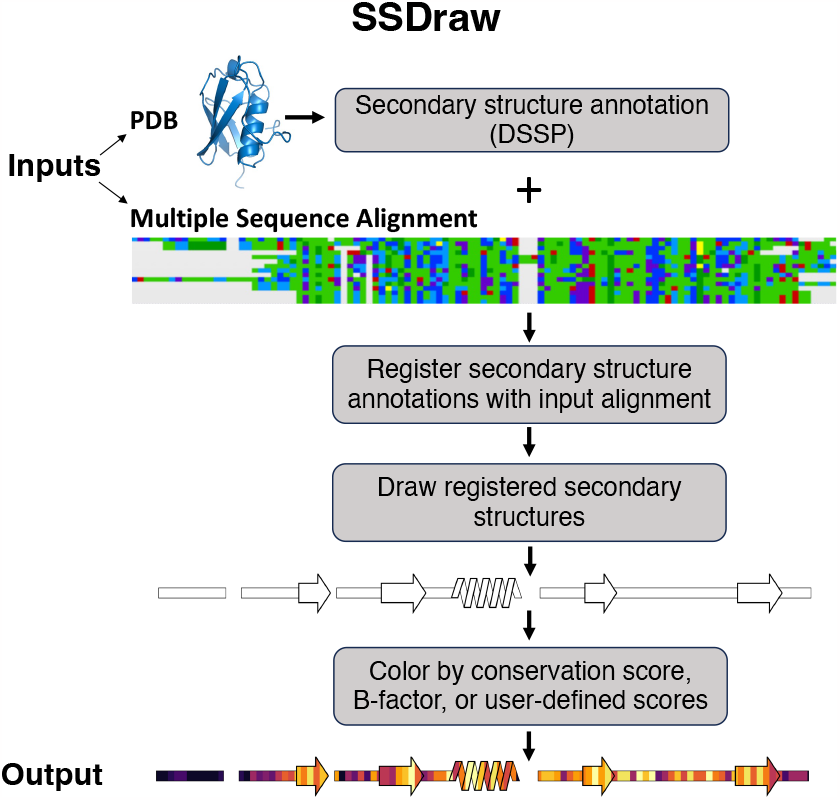
SSDraw fiowchart. Inputs to SSDraw include three-dimensional coordinates of a protein structure in PDB format and a single sequence or multiple sequence alignment in FASTA format. Protein secondary structure is determined from the input PDB using DSSP. Stacked polygons represent α-helices, arrows represent β-sheets, rectangles represent loops, β-turns, disordered regions, and secondary structure shorter than 3-4 residues, and alignment gaps are empty spaces. Secondary structures are registered with the input sequence or alignment to account for gaps and drawn using the Matplotlib patches library for Python3. Finally, secondary structures are colored by sequence conservation scores, B-factor, or another user-defined input. In this figure, the final diagram is colored by default sequence conservation scores. Output structures can be saved as .png, .eps, .svg, .ps, or .tiff files at a user-specified resolution. Multiple sequence alignment depicted using the Alignment Viewer program.

By default, SSDraw computes secondary structure annotations for each amino acid using Define Secondary Structure of Proteins (DSSP)^32;33^, which annotates secondary structure from three-dimensional protein structures based on hydrogen bonding patterns (**Methods**). In lieu of a PDB file, users may input alternative secondary structure annotations^34;35^ or pre-computed DSSP annotations in .horiz format.

Annotated secondary structures are then aligned in register with the input sequence alignment (**Figure 2**) in FASTA format. For proper alignment, the user inputs the name of the reference sequence in the alignment. Protein structures determined by x-ray crystallography or cryo-EM often have unresolved regions due to weak or missing electron density, leading to gaps in their experimentally determined structures. These structural gaps lead to alignment gaps between reference sequences and annotated secondary structures. Accordingly, SSDraw adjusts the reference sequence to be the same length as the secondary structure annotations taken from experimentally determined structures; experimentally unresolved regions are assumed to be disordered and are therefore visualized as loops.

Secondary structures are then drawn with patches from the Matplotlib^36^ package for Python3 (**Figure 2, Methods**). Successive slanted polygons are used to represent α-helices, arrows represent β-sheets, rectangles represent loops, and empty spaces between secondary structures represent alignment gaps. Loops are layered under secondary structures. Segments of regular secondary structure shorter than 4/3 successive residues (α-helices/β-sheets), loops, β-turns, and disordered regions are represented as thin rectangles layered under secondary structure elements (**Methods**).

If desired, secondary structures can be colored by sequence conservation score, B-factor, or another user-defined input (**Figure 1**). This feature was originally developed to compare secondary structure conservation in a family of bacterial response regulators with some secondary structure elements that switch from α-helix to β-sheet in response to stepwise mutation^17^. Sequence conservation scores are computed automatically from the input sequence alignment (**Method**), though scores from Rate4Site^37^, a more accurate conservation metric, may also be inputted. Alternatively, the image can be colored with a solid fill specified by the user. For instance, the first diagram in Figure 2 was generated using a white fill. Custom coloring schemes and custom colormaps may be specified by the user.

If the user wants to assign custom coloring scores to each residue, they have two options. The first is to upload a custom scoring file that contains residue-specific scores. This file is formatted with two columns: column one contains one-letter amino acid codes for each residue to be colored; column two contains scores corresponding to the amino acids in column one; columns are delimited by one space. The second option for custom scoring is to input a PDB file with C-alpha B-factors corresponding to custom scores and coloring the image by B-factor. This option allows the user to easily visualize confidence scores from structure predictors such as AlphaFold2^4^ and ESMfold^6^, if desired. Any range of scores can be used for custom coloring: scores are normalized before the image is colored. Because SSDraw uses the Matplotlib^36^ Python package, any premade Matplotlib colormap may be used; users can also specify custom colormaps as input.

For those desiring to visualize a protein region rather than the whole protein, starting and ending residues can be specified. The Google Colab notebook provides a sliding window that allows the user to select which portion of the alignment will be drawn. Residue numbers corresponding to PDB numbering can be inputted into the local install.

The final output is a linear secondary structure diagram, colored as the user specifies (**Figure 2**). Output files can be saved as .png, .eps, .svg, .ps, and .tiff files at a user-specified resolution. By default, figures are saved as .png files at 600 ppi (pixels per inch), a publication-quality resolution. The user also has the option to include ticks with residue numbers at any specified interval in these final figures.

SSDraw can be used to generate three sorts of outputs: single ungapped diagrams, single gapped diagrams, and multiple aligned and stacked diagrams (**Figure 3**). The first may be best when the user wishes to depict continuous secondary structure of one protein structure (**Figure 3A**), while the second may be preferred if the input structure has unresolved regions. In the latter case, the user may input a sequence with gaps corresponding to unresolved regions in the structure, which will then be depicted as gaps (**Figure 3B**). Finally, the user may wish to generate multiple secondary structure diagrams of homologous proteins for comparison (**Figure 3C**). To accurately compare these diagrams, the secondary structures of each input structure should be aligned to the same MSA. Secondary structures of homologous sequences aligned to different MSAs will likely be unregistered (**Figure 1, top**) and thus cannot be compared. Two examples of more advanced uses of SSDraw are now presented.

**Figure 3:**
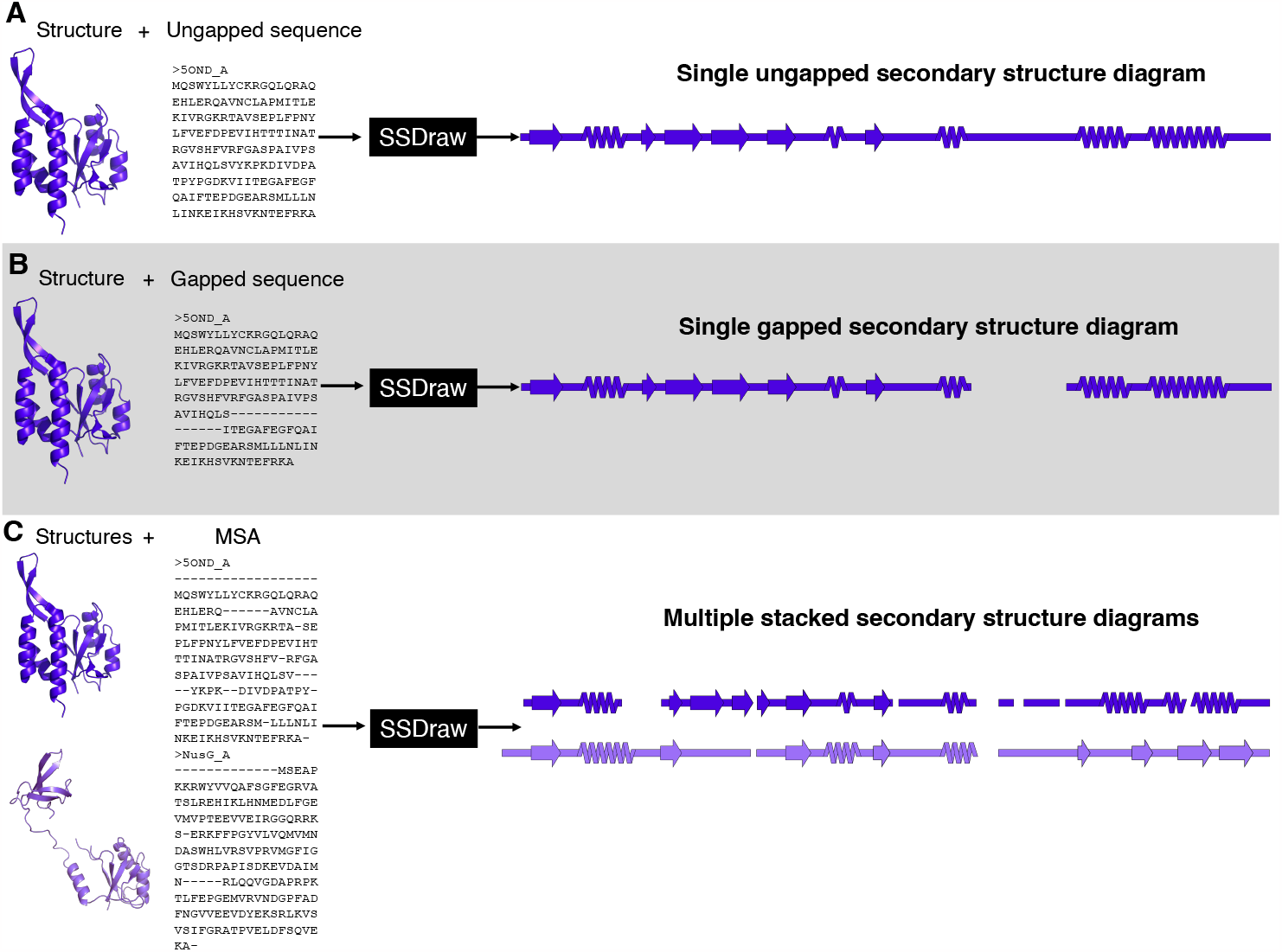
SSDraw has three modes of use. In the first mode, the user inputs a protein structure and an ungapped sequence; a single ungapped secondary structure diagram is outputted (A). In the second mode, the user inputs a protein structure and a gapped sequence; a single gapped secondary structure diagram is outputted (B). In the third mode, the user inputs multiple structures and a multiple sequence alignment that aligns their sequences; multiple stacked secondary structure diagrams are outputted. In all three panels the experimentally determined structure of the transcriptional regulator RfaH^38; 39^ (PDB ID: 5OND, chain A, dark purple) and its sequence (in different alignments) are inputted. In panel C, its homolog NusG^40^ (PDB ID: 6ZTJ, chain CF, light purple) is also inputted. RfaH and NusG are members of the only known family of transcriptional regulators conserved from bacteria to humans^41^. They share a structurally conserved N-terminal domain, while their C-terminal domains differ dramatically in the ground state^42; 43^: RfaH’s is all α-helical, while NusG’s is all β-sheet.

### Advanced example 1: comparing distinct structures with highly identical sequences using a custom color map

SSDraw can be used to compare secondary structures of proteins with high levels of sequence identity but different folds (**Figure 4**). Extensive work has been performed to engineer^15;44-46^ and characterize^16;47;48^ variants of the human serum albumin-binding protein GA and the immunoglobulin binding protein GB. While GA folds into a trihelical bundle, GB folds into a 4β+α structure. One or several mutations can cause the protein to fiip from one ground-state fold to the other^45;46^. The distinct secondary structures of GA and GB variants can be visualized readily with SSDraw (**Figure 4**). The top structure (GB95) is the reference and therefore has no mutations. Three mutations (cyan) to GB95 switch its fold to the three helical bundle (GA95); two mutations to GA95 (magenta) switch it back to the 4β+α fold (GB98). GB98 can be switched back to the trihelical fold with one mutation (yellow), which can be switched back to the 4β+α fold with three mutations (GB98-T25I,L20A, white). Finally, another single mutation (green) switches GB98-T25I,L20A back to the trihelical fold (GB98-T25I). Interestingly, fold-switching mutations tend to occur in the central region of the protein (residues 20, 25 and 30) rather than at the termini, where the closest known fold-switching mutation is 11 residues away from the C-terminus (position 45). Furthermore, all mutations occur in regions of secondary structure rather than loops.

**Figure 4.**
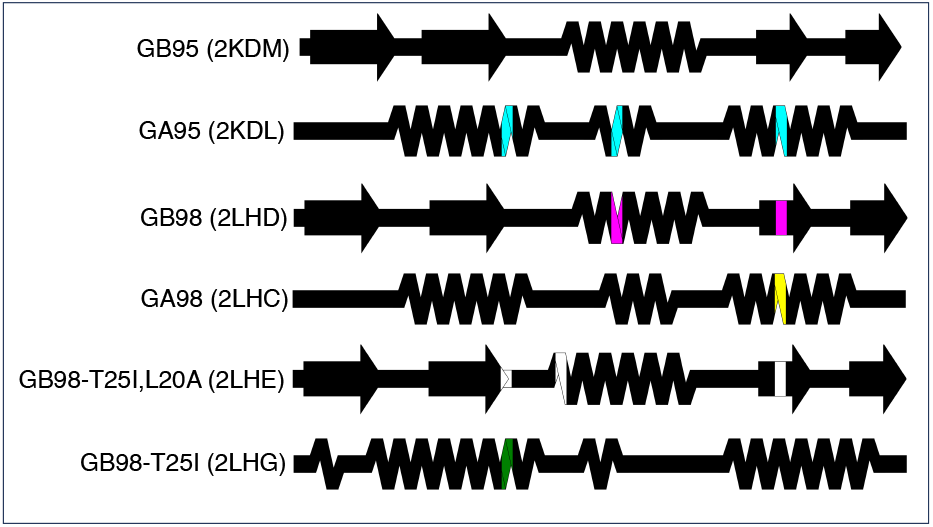
Comparing the structures of proteins with highly identical amino acid sequences but different folds. Diagrams show very different secondary structures derived from the nuclear magnetic resonance structures (PDB IDs in parentheses) of engineered variants of immunoglobulin binding protein GB (4β+α fold) and human serum albumin binding protein GA (trihelical bundle). This figure should be read from top to bottom. Position-specific mutations required to switch a given fold from that of its predecessor, the diagram directly above it, are shown in different colors representing mutations unique to each protein. Black positions were not mutated relative to their immediate predecessors.

### Advanced example 2: comparing sequence conservation in similar structures with a default color map

SSDraw can also be used to relate sequence conservation to secondary structure in protein families with conserved folds. These comparisons for ubiquitin and ubiquitin-like proteins^49^ are shown in **Figure 5**. Not surprisingly, sequences in loop regions tend to be least conserved, while sequences that fold into secondary structures tend to be more conserved. One exception is the second β-sheet, which has been identified as a SUMO1 binding site and putative NEDD8 binding motif by NMR spectroscopy^50^ and structural modeling^51^, respectively. Thus, sequence variation in the second β-sheet may foster different binding functions in different ubiquitin-like proteins. Sequence conservation was calculated directly from the input sequence alignment (**Methods**).

**Figure 5.**
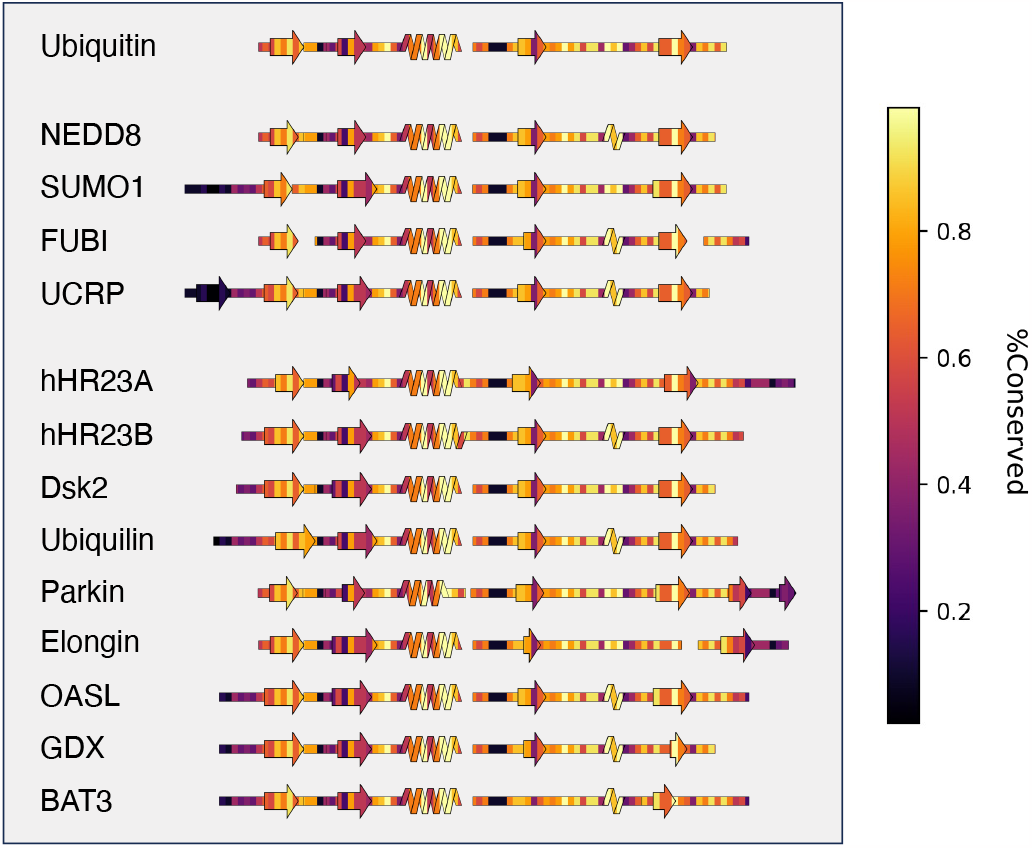
SSDraw diagrams for ubiquitin and ubiquitin-like proteins colored by sequence conservation score (1.0 is highly conserved; 0.0 is least conserved). Sequences of secondary structure elements tend to be more conserved, with the notable exception of the second β-sheet, whose binding functions vary among some ubiquitin-like proteins.

## Discussion and Conclusions

SSDraw generates publication-quality secondary structure diagrams in seconds to minutes. These diagrams can be colored by conservation score, B-factor scores, or a user-specified metric, allowing relationships between secondary structure and other protein properties to be observed readily. SSDraw is expected to be most useful for comparing secondary structures of homologous proteins with different folds, an emerging class of proteins^52^ for which few computational tools are available. Nevertheless, SSDraw may also be used to (1) diagram single structures and color them by any property of interest and (2) compare secondary structures of homologous proteins with conserved folds.

## Methods

### Secondary structure annotation

SSDraw uses DSSP^32;33^ to annotate secondary structure from three-dimensional protein coordinates in PDB format. The local install uses the DSSP module in Biopython^53^ to parse the annotations generated by separate compiled software. Only C-alpha coordinates are necessary for annotation. In addition to regular secondary structure (α-helices and β-sheets), DSSP annotates various local structures such as β-turns and 3_10_ helices. These features are not displayed in SSDraw diagrams because they are not represented well enough. Due to limitations of the patches library, at least 4 consecutive identical annotations (e.g. HHHH or EEEE) would be needed to introduce additional structural elements into these diagrams. **Table 1** shows that α-helices, β-sheets, and loops comprise 87% of all consecutive identical annotations; the next most frequent annotation is Turns, representing 4%. These statistics were calculated from DSSP annotations of 185,725 unique PDB files. Helices are drawn for at least 4 consecutive “H” annotations, and β-sheets are drawn for at least 3 consecutive “E” or “B” annotations, combined in any way. All other annotations are visualized as loops. Short helices with <4 consecutive “H” annotations and short β-sheets with <3 “E” or “B” annotations are also visualized as loops.

**Table 1.** Frequencies of at least 4 consecutive DSSP annotations from 185,725 PDB files.

| DSSP secondary structure annotation | Frequency of at least 4 identical consecutive annotations |
| --- | --- |
| α-helix (H) | 37% |
| β-sheet (E,B) | 36% |
| Loop (‘ ’) | 14% |
| Hydrogen-bonded Turn (T) | 4% |
| $3_{10}$ helix (G) | 3% |
| Bend (S) | 3% |
| 5-helix (I) | 2% |
| Polyproline II helix (P) | 1% |

In some cases, the user who wishes to install SSDraw locally may have difficulty installing DSSP with conda. The user may run SSDraw with the PyDSSP library. PyDSSP is a simplified pytorch-based implementation of DSSP that makes three-state secondary structure annotations (Helix, Sheet, Loop) that match DSSP 97% of the time (https://github.com/ShintaroMinami/PyDSSP).

### Drawing secondary structures

Annotated secondary structures are grouped into three categories: Loop, Helix, and Strand. The lengths of each segment of structure in each category are recorded. Then, each category is drawn separately using the patches library from Matplotlib^36^ for Python3. First, Loops are drawn. Loop lengths are calculated as the number of consecutive annotations divided by 6.0 with the Rectangle patch. When Loops connect elements of secondary structure, they are extended at both ends by 1.0/6.0. All loops have a zorder of 0 so that their images are layered under strand and helix diagrams. Then, coordinates for images of β-sheets and α-helices are stored to be drawn later for better performance. Strands are drawn using the FancyArrow patch with a width of 1.0, linewidth of 0.5, zorder=index increasing over all secondary structures from left to right, head_width of 2.0, and head length of 2.0/6.0. Length is defined as the number of consecutive annotations for the strand being drawn/6.0; to avoid incorrect gapping, this length is extended by 1.0/6.0 if C-terminal elements of secondary structure follow the strand. Helices are drawn as stacked Polygon patches with right-leaning patches layered on top and left-leaning patches layered underneath. The short sides of the polygons measure 1.0/6.0; the long sides measure 1.8/6. Helices begin and end with shorter polygons that align with other secondary structures (height of 1.4/6, width of 1.0/6). All lengths are proportional measures scaled to fit into a figure 25 inches long. Consequently, shorter proteins will have larger secondary structures in the horizontal dimension and vice versa. Vertical heights of all secondary structures are kept constant.

### Coloring secondary structures

Secondary structures have black edges; their insides are filled by clipping an input colormap equal in size to the diagram. Groups of loops, helices, and strands are each converted to clipping paths using Matplotlib’s mpath.Path command. These paths are then converted to patches with mpatch.PathPatch. Finally, an input colormap equal in size to the diagram is generated from user specified parameters or a solid color and clipped to fill the insides of the path (im.set_clip_path command); the rest of the colormap is discarded. Repetitively generating the colormap slows performance considerably. For instance, generating one diagram of a 215-residue response regulator with a mixture of helices and strands (PDB ID: 1A04) takes 1 minute, 5 seconds when a colormap for each secondary structure element–including every polygon to make the helix–must be generated. To improve performance, SSDraw generates colormaps 3 times—once for each class of secondary structure. Running this improved implementation hastened image generation of 1A04 to 2.6 seconds, a 25-fold speed-up from 1 minute, 5 seconds. The Google Colab notebook takes about 2 minutes to generate its first secondary structure diagram because it must load outside software packages, such as DSSP, before running.

### Conservation Scores

Conservation scores are computed directly from an input sequence alignment. First the consensus sequence is determined by calculating the most common amino acids in each column of the alignment. A conservation score is then calculated by:

1. Determining the number, *N*, of amino acids in column *i* with substitution scores ≥ 0 for the consensus residue in column *i*. Substitution scores are calculated using the BLOSUM62^54^ matrix supplied by Biopython^53^.
2. *N* is then normalized by the total number of amino acids in column *i*. Gaps are not included in the normalization.

SSDraw can also take Consurf and Rate4Site scores as input. Consurf scores are taken directly from the input file and used to color the output structure with no modification to the values. Rate4Site scores are normalized and grouped into 9 bins as in^55^.

### Figures

The multiple sequence alignment in Figure 2 was generated using Alignment Viewer: https://github.com/sanderlab/alignmentviewer. All three-dimensional protein structures in Figures 2 and 3 were generated using PyMOL.^56^

## Acknowlegements

We thank Myeongsang Lee and Joseph Schafer for testing local installs of SSDraw and Leslie Ronish and Joseph Thole for testing the Google Colab notebook. L.L.P. thanks Loren Looger for inspiring the idea of SSDraw. This work was supported by the Intramural Research Program of the National Library of Medicine, National Institutes of Health (LM202011, L.L.P.).

## Code availability

The complete code, documentation, and examples for SSDraw can be found at: https://github.com/ethanchen1301/SSDraw. A Google Colab notebook is also avilable at: https://colab.research.google.com/github/ethanchen1301/SSDraw/blob/main/SSDraw.ipynb. To upload local files into the Colab notebook, the user must run the notebook with Google Chrome.

